# Validated Bayesian differentiation of causative and passenger mutations

**DOI:** 10.1101/097931

**Authors:** Frederick R. Cross, Michal Breker, Kristi Lieberman

## Abstract

In many contexts, the problem arises of determining which of many candidate mutations is the most likely to be causative for some phenotype. It is desirable to have a way to evaluate this probability that relies as little as possible on previous knowledge, to avoid bias against discovering new genes or functions. We are isolating mutants with blocked cell cycle progression in *Chlamydomonas*, and determining mutant genome sequences. Due to the intensity of UV mutagenesis required for efficient mutant collection, the mutants contain multiple mutations altering coding sequence. To provide a quantitative estimate of probability that each individual mutation in a given mutant is the causative one, we develop a Bayesian approach. The approach employs four independent indicators: sequence conservation of the mutated coding sequence with *Arabidopsis*; severity of the mutation relative to *Chlamydomonas* wild type based on Blosum62 scores; meiotic mapping information for location of the causative mutation relative to known molecular markers; and, for a subset of mutants, transcriptional profile of the candidate wild type genes through the mitotic cell cycle.

These indicators are statistically independent, and so can be combined quantitatively into a single probability calculation. We validate this calculation: recently isolated mutations that were not in the training set for developing the indicators, with high calculated probability of causality, are confirmed in every case by additional genetic data to indeed be causative. Analysis of best reciprocal blast relationships among *Chlamydomonas* and other eukaryotes indicate that the Ts-lethal mutants that our procedure recovers are highly enriched for fundamental cell-essential functions conserved broadly across plants and other eukaryotes, accounting for the high information content of sequence alignment to *Arabidopsis*.

## Introduction

The use of model genetic systems to obtain insight into related organisms is well established; as a leading example, yeast genetics has been highly revelatory about fundamental cell biology in animals (Botstein and Fink 2011). Because fungi diverged from animals considerably after their last common ancestor diverged from the plant lineage (Rogozin et al., 2009), it is an open question to what extent yeast/animal paradigms will apply to the plant lineage.

To address this question, we initiated a genetic screen for Ts-lethal mutations in the green alga *Chlamydomonas reinhardtii*, focusing on cell cycle control mutations (Tulin and Cross 2014). The reasoning was that facile microbial genetics and cell culture would facilitate isolation of informative mutants, compared to carrying out a related screen directly in higher plants. A specific feature likely to make such a screen easier in green algae than in higher plants is the high degree of gene duplication in plants, due largely to multiple whole-genome duplications in the plant lineage after divergence from green algae. Loss-of-function genetics is severely hampered by the presence of duplicated sequences; the *Chlamydomonas* genome is largely (though not entirely) single-copy for protein coding sequence.

From the broad spectrum of Ts-lethal phenotypes, we concentrated on two classes: mutants that initiated some cytological features of the cell division program but failed to complete division, and mutants with competence at cell growth, that were unable to initiate division processes. We called these mutant classes ‘*DIV*’ and ‘*GEX*’ respectively. Obviously cell-cycle-related genes based on annotations (e.g., DNA polymerase subunits) were mainly in the *DIV* class; *GEX* genes were more diverse in function based on annotations. Together, these classes represented a few percent of the total Ts-lethal spectrum (Tulin and Cross 2014).

The assumption behind the project, with respect to gaining insight into higher plant genomes, is that evolutionary conservation will likely preserve function of essential genes, so that genes identified by Ts-lethal screening as essential in *Chlamydomonas* will be essential (or will be members of essential sequence families) in higher plants. This assumption is plausible but by no means certain. As a trivial example, yeast cell walls are essential, but animal cells lack walls and genes for their production. Additionally, replacement of important cell cycle regulators by entirely unrelated proteins carrying out similar functions is documented comparing yeast and animals (Cross et al., 2011, Medina et al., 2016).

In our approach, Ts-lethal mutations were induced by UV mutagenesis; identification of likely causative mutations was by next-generation sequencing analysis of bulked segregant pools, which identifies SNPs at or linked to the causative mutation (Tulin and Cross 2014). Two problems remain. First, the sequencing approach identifies the causative mutation in most but not all cases; second, the density of UV-induced mutagenesis needed for efficient screening (Breker et al. 2016) is such that the causative SNP is frequently very difficult to separate by meiotic recombination from a small number of linked ‘passenger’ SNPs. These problems mean that formally, in no case can we assert with certainty that a given SNP is causative – if it is one of a number of linked candidates, *a priori* any of them could be causative, and even if it is the only candidate, it is possible that it is a passenger with an unsequenced, truly causative mutation.

To solve this problem, we followed three routes. First, complementation and linkage analysis can demonstrate that we have multiple independent alleles in the same gene. In such a case, if independent mutations altering the same gene model are found by sequence analysis, we assume that these mutations are causative. Second, in many cases we could isolate revertants by selection at high temperature. When sequence analysis showed that the revertants altered one of the gene models hit by the original set of SNPs (typically by exact reversion, or by pseudo-reversion at a nearby residue), we assume that this also constitutes definitive identification.

Third, for mutants not covered by either method, we developed a Bayesian method, based on sequence characteristics of the candidate SNPs and linkage analysis. This method was sketched out previously (Tulin and Cross 2014). Here, we specify and develop the exact model. We extend the method, substantially increasing its power, by addition of a new indicator: transcriptional profiling of candidate genes through the cell cycle. Critically, we validate the method by analysis of new genetic and molecular data, with mutations not in the training set used to develop the indicators. Finally, we use bioinformatics tests, including analysis of best reciprocal Blast-defined sequence families, to show that essential genes in *Chlamydomonas* are preferentially conserved with respect to higher plant genomes.

## Methods

### Mutant isolation and mutation detection

Temperature-sensitive-lethal (‘Ts−‘) mutants were isolated after UV mutagenesis, and screened for proliferation-specific defects as described (Tulin and Cross 2014; Breker et al., 2016). Candidate causative mutations were identified by bulked segregant sequence analysis, as described (Tulin and Cross 2014), or by a new method for parallel bulked segregant sequence analysis in combinatorial pools (Breker et al., in preparation). Candidate causative mutations were those that were uniformly mutant in pools of Ts− segregants from a cross to wild type (or, reciprocally, uniformly wild type in pools of Ts+ segregants.

We call a candidate causative mutation ‘definitive’ (Tulin and Cross 2014) if it hits a gene model that is also hit in an independent isolate, and the two isolates fall in the same linkage/complementation group; or if we have isolated an intragenic revertant that alters the mutation (true or pseudorevertants). For the present analysis, we employed definitive mutations defined in Tulin and Cross (2014). ‘Passenger’ mutations (UV-induced mutations that do not cause the Ts− phenotype) were identified either as those present at <100% prevalence in pools of Ts− segregants from crosses of various mutants to wild type, or that were uniform but distinct from the true causative mutation, provided the latter was known ‘definitively’. In this way we collected 67 independent definitive causative mutation, described in Tulin and Cross (2014), and 137 passenger mutations. These mutations constitute the ‘training set’ for finding Bayesian discriminators.

Importantly for development of the present analysis, identification of the causative and passenger mutations in the training set is entirely based on genetic data, and is independent of annotations (beyond the essential segmentation of the genome into gene models [Blaby et al. 2011]), mutational severity or transcriptional pattern.

For the ‘test set’, we employed the 20 mutants isolated in Tulin and Cross (2014) as single members of their complementation groups. All of these mutants were mapped to a chromosomal location relative to physical markers (generally to within 1-2 Mb), and all had varying numbers of candidate mutations for causality within the mapped region. For seven of these mutants, additional mutant screening (Breker et al., 2016) yielded new alleles based on linkage and complementation testing, and genome sequences were obtained for these new mutants as well.

### Linkage mapping

We carried out meiotic mapping of Ts-lethal mainly employing two methods. We develop allele-discriminatory PCR probes (using the competitive approach described by Onishi et al. 2016) that allow determination of WT or mutant in a single reaction, and test multiple meiotic segregants (Ts− or Ts+) for marker status. This procedure has the benefit of anchoring the mapping at exactly the site of interest, but the disadvantage that it is hard to test large numbers of progeny. Ts− segregants lacking the mutant marker, or Ts+ segregants with the marker are counted as recombinant chromosomes. A second method is tetrad analysis of a mutant against a tester ts-lethal with a known physical location. We have developed a ‘micro-tetrad’ approach in which hundreds of tetrads can be rapidly dissected and analyzed on a single plate, by dissecting over an area of ~1 mm and transferring the plate to restrictive temperature only when meiotic products have just germinated and formed microcolonies. It is then simple to count the number of Ts+ segregants per tetrad to determine PD, NPD and T tetrad classes.

For analysis, we combine mapping data obtained with independent alleles of the same gene, defined by non-complementation and lack of recombination in meiosis, since the latter property, using our current recombination assay (Breker et al., 2016) implies no recombination in hundreds or thousands of meiosis, thus very close linkage.

The computation of probability requires specification of a physical region that is assumed to contain the causative mutation. For simplicity, we use as boundaries positions 2 Mb away from the leftmost or rightmost SNPs detected as uniformly mutant in bulked-segregant sequence analysis.

### ‘Best Reciprocal Blast’ analysis

If two sequences, one from each of two genomes, find each other as their best BLAST hit in cross-genome comparisons, this provides the basis of forming a potential ‘orthologous’ family, containing these two sequences as well as potential recent gene duplicates from within each genome (‘in-paralogs’) (Remm et al., 2001). We wrote MATLAB code to carry out essentially this procedure (code provided in S.I.). Here we report analysis of such orthologous families ‘seeded’ by search with each *Chlamydomonas* protein as query against seven genomes: *Arabidopsis*, *Brachypodium*, *Physcomitrella* (dicot, monocot, and moss from the land plants); *Homo* and *Drosphila* (two animals); *Saccharomyces* and *Aspergillus* (two fungi). Reference proteomes were downloaded from public-access databases Phytozome, UniProt, SGDB and ADB.

## Results

### The training set

To determine useful Bayesian indicators, it is necessary to have a training set of positive and negative examples. To obtain these, we made the assumption that causative mutations were ‘definitively’ (i.e., ground-truth) detected in a specific gene model in the following cases: (1) where multiple alleles in the same gene, defined by failure of genetic complementation and recombination, were shown to have mutational lesions altering coding sequence in the same gene model; (2) where at least a subset of selected revertants of the mutant were found to have reverted the original lesion in the gene model (either exact or pseudo-revertants). In almost all cases, these assignments were also supported by meiotic mapping. These constitute positive examples of causative mutations. The bulked-segregant sequence analysis to determine candidate causative mutations (Tulin and Cross 2014) also yields sequence of non-causative mutations induced by UV (‘passengers’). We consider such mutations to be definitive passengers either if they are either linked to but distinct from a definitive causative mutation, or if they are on a different chromosome from the causative mutation. (Note that we usually can definitively identify the chromosome bearing the causative mutation even when the causative mutation itself is not definitively identified, based on uniform detection in the region of mutations [frequently passengers] linked to the causative mutation). This analysis resulted in a training set of 69 causative and 137 passenger mutations.

The training set is only useful if it is an unbiased sample. In this case, bias appears unlikely. Identification of the causative mutations is based entirely on formal genetic criteria, and is independent of any annotation information (e.g., alignment, previous information about the gene or its relatives).

### BLAST-detected conservation with *Arabidopsis*

The *DIV/GEX* ts-lethal mutations identify essential functions, and genes required for essential functions might be more conserved across evolution than non-essential genes. The land plant *Arabidopsis* and *Chlamydomonas* diverged ~1 billion years ago (Yoon et al 2004), and their common ancestor diverged from other eukaryotes including animals and yeast about 1.6 billion years ago. Only 40% of *Chlamydomonas* genes have detectable BLAST homology over any part of their sequence with *Arabidopsis*, and this number drops to 30% with a requirement for a BLAST bit-score of at least 100 (data not shown). Thus conservation with *Arabidopsis* is a strict criterion, but one that likely allows detection of core machinery conserved in the Viridiplantae lineage.

Retaining protein function through evolution generally requires much greater conservation in some protein regions (e.g., active sites) than others (e.g., protein loops, N- and C-termini). The scheme used here (Figure 1) subdivides the relationship of mutational position to BLAST results as follows:

**Figure 1.**
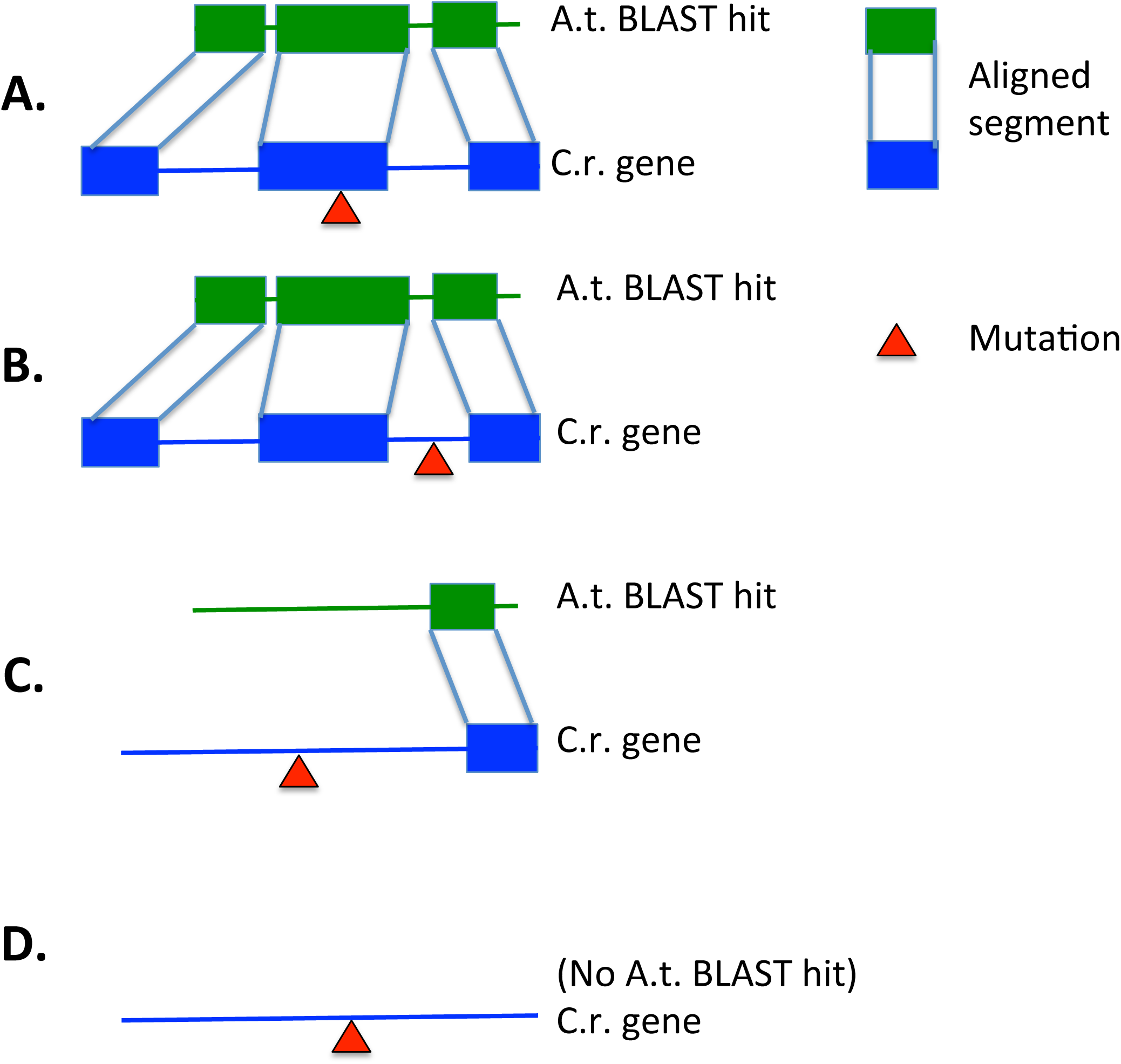
Relationship of mutation to BLAST alignments of *Chlamydomonas* gene model to *Arabidopsis*. A: mutation falls within a conserved segment; B: mutation falls between conserved segments; C: mutation is N- or C-terminal to a conserved segment; D: no conserved segment detected by BLAST. Note that for empirical classification, a mutation in an unconserved residue, in an unaligned residue within a high-scoring pair (conserved segment), or a mutation falling between two conserved segments are all assigned to class B. Rules for truncating mutations (stop codons, splice donor/acceptor mutations; found in a small minority of the causative Ts-lethal mutations): if upstream of the C-terminal-most conserved segment, class A; otherwise class B.

Class A: mutation falls within a segment of BLAST alignment (high-scoring pair or HSP), and the mutation alters a conserved residue within this segment. [‘Conserved’ is defined operationally as follows: the Blosum62 score (Henikoff and Henikoff 1993) between *Arabidopsis* and *Chlamydomonas* at the position is greater than 0 (meaning that conservation is observed more often than would be expected by chance).]

Class B: mutation falls within an overall conserved region, but alters an unconserved residue; is BLAST-aligned across a small deletion in the *Arabidopsis* sequence in the HSP; or is found between two distinct HSPs. (These three distinct possibilities are joined into one category to prevent excessive slicing of the training set, and because in all cases the mutation is to an unconserved residue but is surrounded by regions of conservation).

Class C: mutation is N-terminal or C-terminal to all detected HSPs.

Class D: no *Arabidopsis* Blast hit.

The order A>B>C>D seems plausible for likely disruption of conserved protein structure/function. While finer-grained classifications are possible, it is important to keep the number small enough that the training set can provide reasonable occupancy of the bins.

The distributions of causative and passenger mutations from the training sets among these classes showed sharp differentiation (Figure 2, top left). Most notable was the extreme enrichment of classes A and B among causative mutations, indicating a strong correlation between essential functional regions of key proteins, and conserved alignment through evolution.

**Figure 2.**
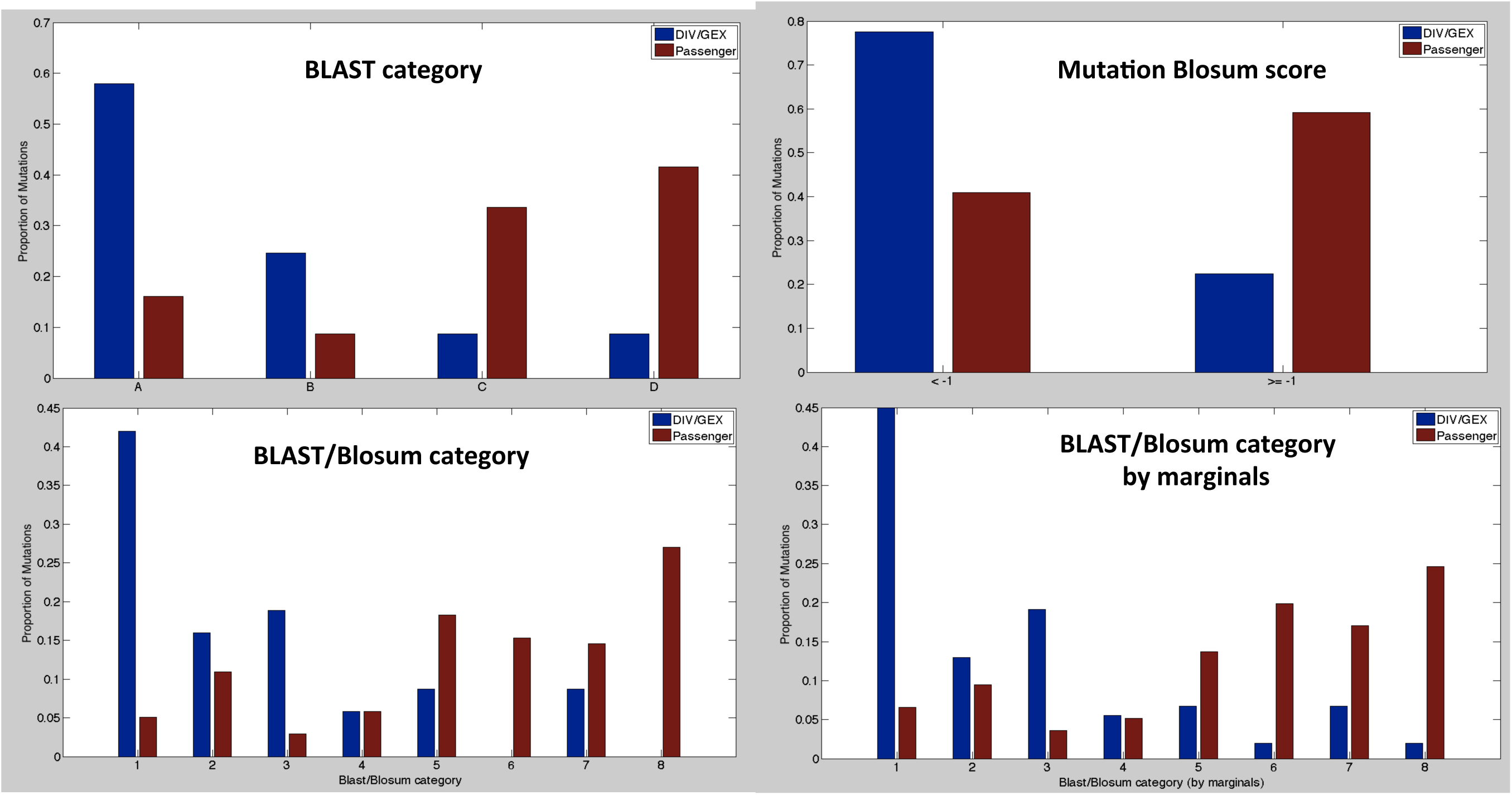
Constructing Bayesian classifiers. Mutational Blosum scores, BLAST category, and Blast/Blosum category distribution (top left, top right, bottom left) for definitive DIV/GEX vs. coding-sequence-changing passenger mutations (the training set). Lower right: Blast/Blosum categories computed by multiplication of individual Blosum and Blast probabilities. Near identity of the two lower graphs indicates independence of these measures.

### ‘Best Reciprocal Blast’ analysis

Blast alignments can reflect a range of degree of relationship. High, end-to-end similarity might reflect orthologous function. However, many alignments are due solely to conservation of a common small protein domain (e.g., WD40-repeats). One way to discriminate among Blast hits is detection of ‘best reciprocal Blast’ (‘BRB’) hits: when protein X from genome A finds protein Y from genome B as one of the best hits, and reciprocally, protein Y from genome B has protein X from genome A as one of the best hits (Remm et al., 2001). Frequently there are other sequences in genome A, more similar to X than is protein Y from genome B; these may be derived from duplication of X after separation of lineage A from lineage B (Figure S1). We have carried out reciprocal Blast analysis along these lines (Matlab code in S.I.) between *Chlamydomonas* and a range of other eukaryotes: three representative land plants: *Arabidopsis*, a dicot; *Brachypodium*, a monocot; *Physcomitrella*, a moss; two animals, humans and *Drosophila*; two fungi, *Aspergillus* and *Saccharomyces*.

To validate the BRB search, we note that *Chlamydomonas* should be approximately equally diverged from *Arabidopsis*, a dicot, *Brachypodium*, a monocot; *Physcomitrella*, a moss; since these are all in the higher land plant lineage with a single divergence time from *Chlamydomonas*. Consistent with this idea, a Venn diagram shows very strong overlap between *Chlamydomonas* proteins identified as BRB family members using these three genomes as targets (Figure 3, top left). This set of 2818 orthologous families may be broadly distributed among Viridiplantae.

**Figure 3.**
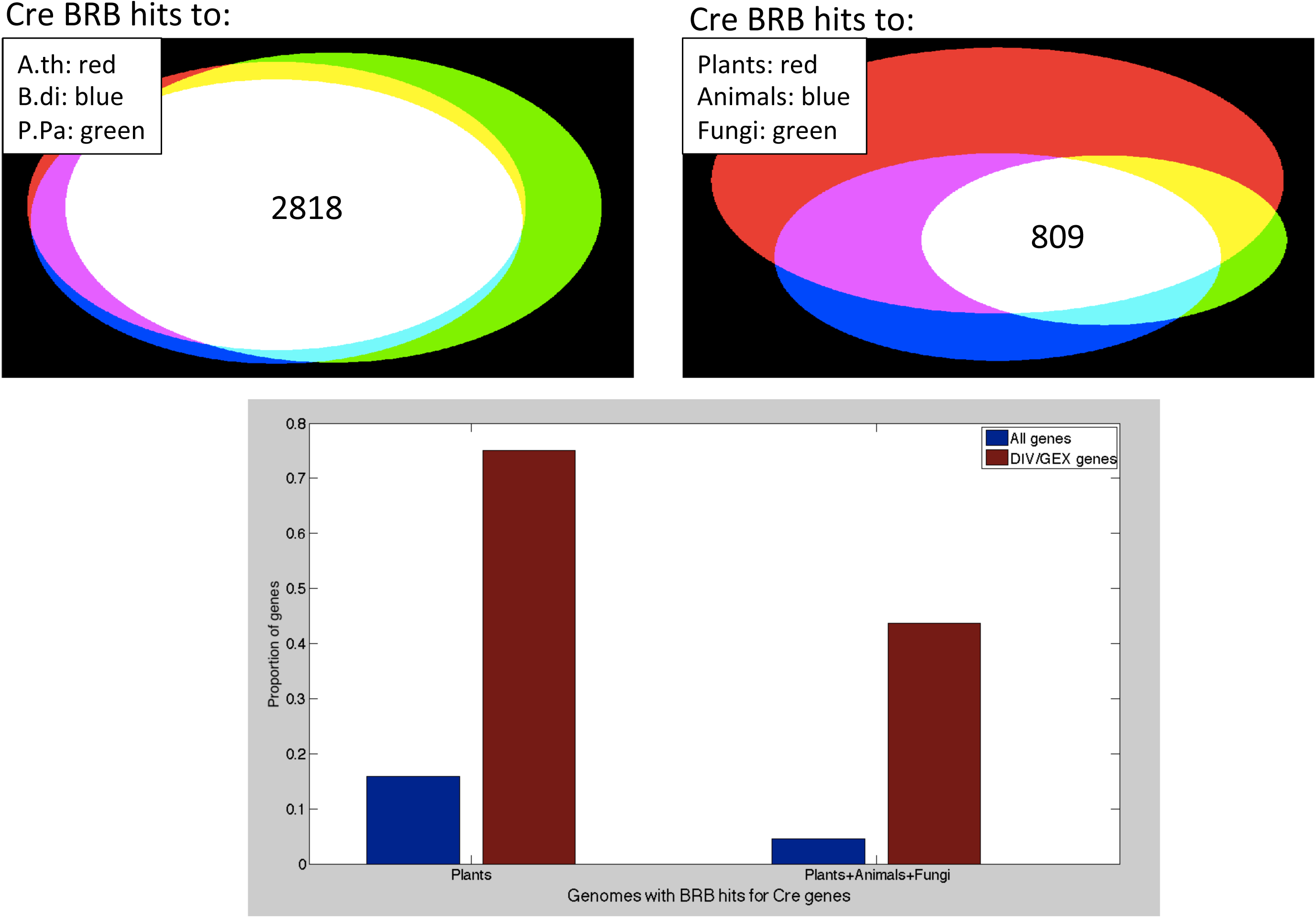
Best-reciprocal BLAST analysis was carried out with the *Chlamydomonas* proteome as query, against *Arabidopsis thaliana*, *Brachypodium distachyon*, *Physcomitrella patens*, *Homo sapiens*, *Drosophila melanogaster*, *Saccharomyces cerevisiae*, *Aspergillus niger* genomes (three plants, 2 animals, 2 fungi). Top left: overlap of identity of *Chlamydomonas* genes in BRB-orthologous families with the three plant genomes. 2818 *Chlamydomonas* genes are in such families with proteins from all three plant proteomes. Top right: overlap of *Chlamydomonas* genes in BRB-orthologous families with all three plant proteomes, both animal and both fungal genomes. Below: proportion of total *Chlamydomonas* genes (blue) and definitive DIV/GEX genes (red; Tulin and Cross 2014) in the overlap classes shown at top.

In the *Chlamydomonas* vs. *Arabidopsis* search, most candidate ortholog families contain single members in *Chlamydomonas*, but frequently contain multiple members in *Arabidopsis* (Figure 4). This may be due to ancient gene or genome duplications in the land plant lineage (Adams and Wendel 2005), after the split from green algae. Similar results for gene family sizes were obtained in comparisons of *Chlamydomonas* to *Brachypodium* and *Physcomitrella* (data not shown).

**Figure 4.**
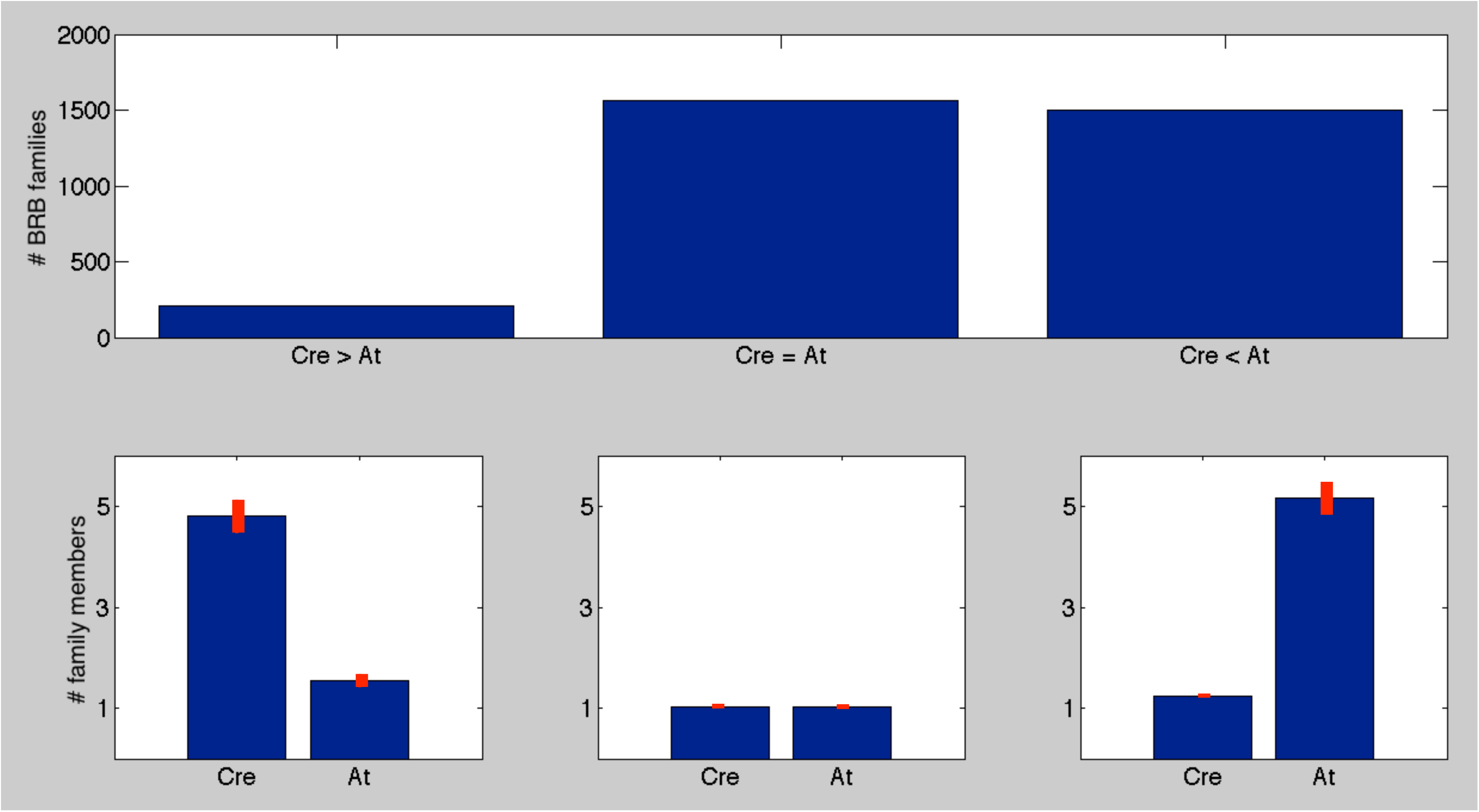
Gene duplicates in BRB-defined orthologous families. Orthologous families containing *Chlamydomonas* (Cre) and *Arabidopsis* (Ath) members were sorted by which proteome contributed more members to the family (top). Left: Cre>At; middle: same numbers from each; right: At>Cre. Bottom: mean and SEM number of family members from each proteome, in each case.

809 *Chlamydomonas* genes were in orthologous families with family members in all seven genomes tested (three plant, two animal, two fungal); these may be near universal among eukaryotes.

### *DIV/GEX* gene enrichment in the BRB gene sets

In carrying out this mutant hunt (Tulin and Cross 2014), the phenotypic sorting of mutants, and concentration on the ‘*DIV/GEX*’ class, was based on the idea that genes with these mutant phenotypes might specifically identify core conserved cellular functions. Confirming this idea, while only 26% of *Chlamydomonas* genes are in a BRB-defined orthologous family with any of the seven genomes tested, 79% of *DIV* and *GEX* genes are in a BRB family with a gene from at least one of the test genomes. This enrichment is even greater in the Plant-orthologous families (76% vs. 16%), and is especially strong in the Plant-Animal-Fungi orthologous families: (44% vs. 5%) (Figure 3). This finding strongly suggests that the *DIV/GEX* class is strongly enriched for deeply conserved functions, dating back to the LCA of plants and Opisthokonts (thus near to the eukaryotic LCA [Rogozin et al., 2009]).

### Mutational severity

Amino acid substitutions vary in their potential to disrupt protein function. To measure the likely severity of effect of substitutions caused by UV-induced mutations, we use the Blosum62 score (Henikoff and Henikoff 1993). Blosum scores have been shown to perform well compared to most other measures for determination of mutational severity (Yampolsky and Stoltzfus 2005). Distributions of this score for causative and passenger mutations are broad, but there is a clear shift of the causative mutations to more negative (more severe) Blosum62 scores. Empirically, a cutoff of score <-1 gives a good separator between most causative and most passenger mutations (Figure 2, top right).

### ‘Blast/Blosum’ combined index

To make a simple discriminator, the Blast classes AD were combined with the severity index (mutation has Blosum <−1, or >=−1) to make eight classes. Notably, the occupancy of these classes by causative and passenger mutations was almost exactly that expected from multiplication of the marginals – that is, the Blast criterion and the mutational severity criterion were almost entirely independent (compare Figure 2 lower left and lower right). The overall discriminatory power of this index is high, with log10-likelihood ratios ranging from 0.9 for category 1 to −1.4 for category 8 – that is, up to a 200-fold differential.

The distribution of passenger mutations among these categories was similar to that of mutations randomly generated *in silico* (data not shown), suggesting that the passenger training set is largely reflective of the initially generated mutational spectrum, with little effective selection operating against most mutations over the short term of these experiments.

### Construction of a formal Bayesian model

The full presentation is in the Appendix. Briefly, suppose there are N candidate SNPs. We assume the causative mutation is a single-gene lesion (since a large majority of Ts-lethals segregate 2:2 in tetrad analysis). Note that in the present experimental context, N is generally not large, since the bulked segregant sequencing approach (Tulin and Cross 2014) will rule out most of the mutations in the original mutant since the WT and mutant alleles will both be detected (typically around 50% each) in sequence from a pool of ~10 ts segregants from a cross to WT. This reduces N from hundreds to single digits.

Bulked segregant sequencing fails to detect *any* candidate causative SNP in a minority of cases (Tulin and Cross 2014) (discussed further below). The reason for this is presently unknown; a simple explanation would if the mutated gene is not present in the assembled genome (which is known to have gaps), but this is unlikely to be the complete explanation (Tulin and Cross 2016). Call U the probability (which we estimate at ~~25% [Tulin and Cross, 2014]) that the causative mutation escaped detection by sequencing and therefore is none of the N candidates. If the causative mutation was detected by sequencing, we assume it is exactly one of the N SNPs.

From Figure 2, define B_i_ as the likelihood ratio for SNP_i_: the probability that a causative mutation is in the Blast/Blosum class of the SNP, divided by the probability that passenger mutation is in this class. Define Q as the likelihood ratio of unsequenceability U/(1-U).

Then the probability that one specific SNP_i_ is causative, given the Blast/Blosum characteristics of the collection of N SNPs, by Bayes’ theorem (see S.I. for detailed derivation), is:

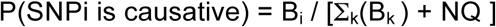

The probability that the causative mutation escaped detection by sequencing is:

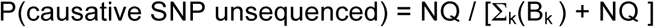

Mutations with high B_i_ (such as severe mutations in conserved residues) are more likely to be causative. Increasing numbers of candidate mutations (higher N) decreases likelihood that any individual one is causative. Conversely, the likelihood that the causative mutation escaped detection by sequencing decreases with increasing numbers of SNPs with high B_i_.

### Mapping information

In principle, meiotic mapping could reduce the interval carrying the causative mutation to arbitrarily small size. However, meiotic distances are in centiMorgans (cM). Conversion to physical distance (needed for causative SNP identification) requires knowing the cM/megabase (Mb) conversion ratio. This is known on average to be ~10 cM/Mb (Merchant 2007; Tulin and Cross 2014), but it is well known in many organisms that this ratio is variable across the genome due to hot- and cold-spots for recombination. We have found one region of ~0.5 Mb across which there is no detectable recombination (no recombinants in at least 1000 meioses; unpublished results); this is an extreme case, but surely not the sole such interval.

However, in case where candidate mutations are separated by 1 Mb or greater, meiotic mapping can provide discriminatory power, using the mapping strategies described in Methods. It is useful to translate such mapping results into estimated probabilities for physical location of the causative mutation. The full development of these estimates is described in Supplementary Information. Briefly, we assume a normal-like distribution of probable location of the causative mutation, with the mean at the best estimate (measured cM from the known marker * 0.1 Mb/cM), and standard deviation 0.5 Mb (based on the maximum non-recombining interval detected, and on apparent measured error (deviation from the 0.1 Mb/cM average) in many such mapping experiments [Tulin and Cross, 2014]). Passenger mutations have probability density that is uniform across the known possible region (typically a chromosome or chromosome arm). Then if L_i_ is the location of SNP_i_, g(L_i_) is a scaled relative probability density at this position, and Q is the likelihood ratio that the causative mutation was not sequenced (that is, if U is the probability of unsequenceability, Q=(U/(1-U)), then the probability that of N SNPs, SNP_i_ is causative is:

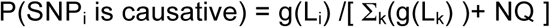

- where L_i_ is the location of SNPi, and g is an appropriately scaled probability density function intended to conservatively represent plausible locations of the causative SNP.

(Derivation of g (S.I.) essentially assumes that a given mapping result is equivalent to an approximately normal distribution for probability of true location, with mean the best estimated location, and standard deviation based on mapping error).

Chromosomal location is almost surely independent of Blast/Blosum values, since like most eukaryotes, *Chlamydomonas* exhibits broad dispersal of functionally related genes across chromosomes. Independent probabilities multiply; therefore, Blast/Blosum information can be integrated with mapping information to produce a single probability of causality for each candidate SNP:

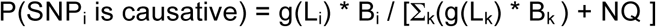

### Transcriptional regulation

*Chlamydomonas* exhibits very strong differential transcription through its mitotic cycle. In particular, many genes, including many that are probably specifically required for DNA replication and cell division, are induced by huge factors (>100-fold) in S/M-phase cells compared to newborn G1 cells (Tulin and Cross, 2015; Zones et al., 2015). The mutations we isolated previously as blocking cell cycle progression (Tulin and Cross 2014) were separated into two broad phenotypic categories: ‘*div*’ mutations, that showed evidence of entry into the replicative cycle followed by arrest, and ‘*gex*’ mutants, that showed no signs of even initial replication/division processes. It was noted (Tulin and Cross, 2015; Zones et al., 2015) that *DIV* genes, but not *GEX* genes, were highly likely to exhibit the transcriptional pattern noted above, of huge induction in S/M phase. Since this induction is observed with only a small proportion of genes overall, this provides another plausible Bayesian discriminator, specifically for the *DIV* class of genes.

To set up such a discriminator, we made use of cell-cycle transcriptome derived from RNAseq of two replicates of a light-dark-synchronized cell cycle (Zones et al., 2015). We converted data for every gene into two numbers: peak-to-trough ratio (PTR) and time of peak (T) (Figure 5A). Almost 80% of genes are at least 2-fold differentially regulated through the timecourse, as reported (Zones et al., 2015). It is also notable that peak times are mostly in three clusters: early (around 6 hrs, before commitment to a replicative cell cycle [Zones et al., 2015), middle (around 13 hrs, in the middle of the S/M cycles); and late (20-24 hrs, as newborn cells mature and hatch). *DIV* genes identified in Tulin and Cross (2014) are marked on the figure. Most *DIV* genes are expressed in the S/M period, at very high PTR, clearly separated from the large majority of other genes (Zones et al., 2015, Tulin and Cross 2015).

**Figure 5.**
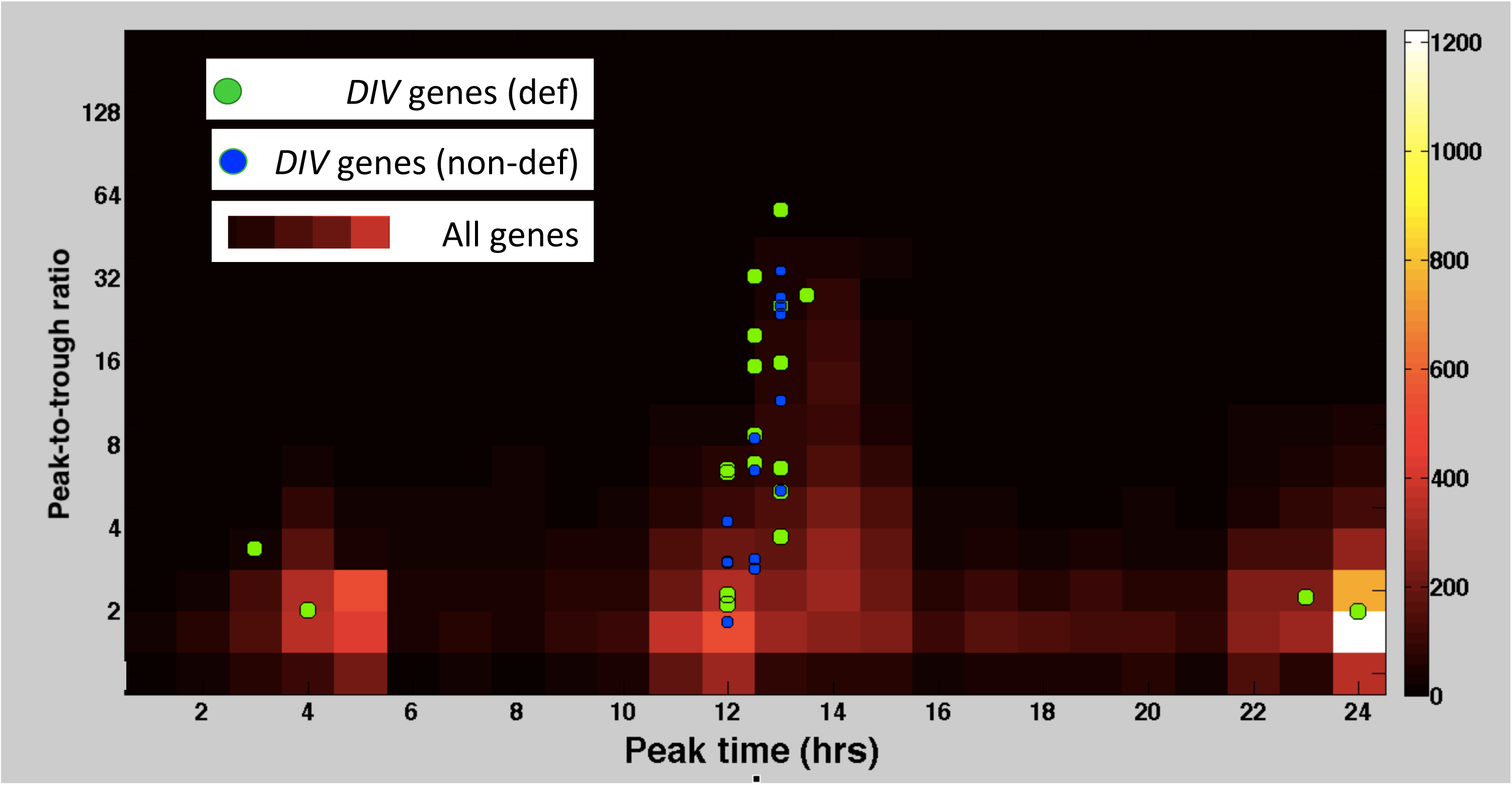
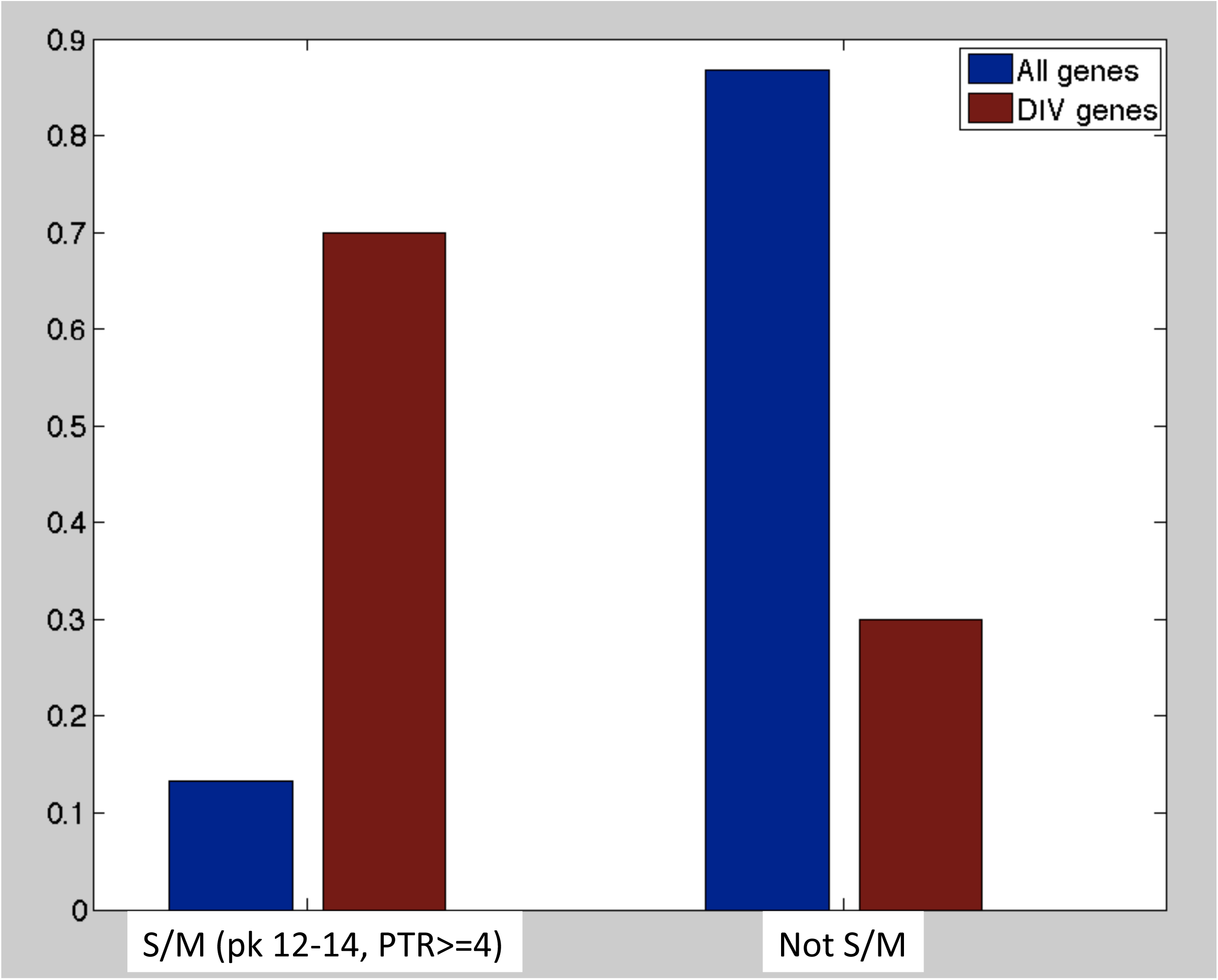
The *DIV-like* transcriptional pattern. A. Cell cycle transcriptional data of Zones et al. (2015) was analyzed: two timecourses were averaged, and a three-timepoint running average calculated. Two values were extracted: peak time (time point at which the plot was maximal), x-axis; and peak-to-trough ratio (highest level divided by lowest level), y-axis. The heat map (scale at right) shows the complete gene set. Placement of ‘definitive’ and ‘non-definitive’ *DIV* genes on the plot is indicated by green and blue circles. B: proportion of *DIV* genes and of all genes in two bins of PTR/peak expression time, the ‘S/M’-like pattern (PTR≥4, peak time 12-14 hrs; Zones et al., 2015), and all other patterns.

The number of definitively identified *DIV* genes is not large, so the data will not at present support a fine-grained analysis. ~70% of *DIV* genes have T between 12 and 13.5 hrs, and PTR>=2, while this category includes only 13% of the total genes. We used this criterion as a binary discriminator to detect a ‘*DIV*-like’ pattern (Figure 5B). Importantly, this criterion was established using only the ‘definitively’ identified *DIV* genes from Tulin and Cross (2014), although it is evident that the broader class of probable *DIV* genes follows the same pattern (Figure 5A). This criterion provides another Bayesian test. Call ‘T_i_’ the transcriptional likelihood ratio (*DIV*/all genes) for the gene in which SNP_i_ is found (two classes, ‘*DIV*-like’ or not):

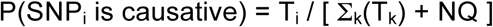

Cell-cycle transcriptional pattern is largely independent of BLAST homology to *Arabidopsis* (Figure S3), and is surely independent of mutational Blosum scores for individual SNPs. Absence of functional clustering implies transcriptional pattern is likely also independent of map position. Therefore, the test for transcriptional category can be combined multiplicatively with Blast/Blosum scores and mapping information to yield an integrated probability that a given SNP is causative for a mutation in the *div* phenotypic class:

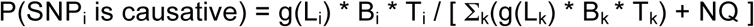

The transcriptional regulation test is more preliminary than the Blast/Blosum and mapping tests, because it is based on only a small training set, and is restricted to a single phenotypic class. This will doubtless improve as more mutants are defined, growing the training sets for *div* and for other phenotypic classes. Initial analysis supported some subclustering among the *gex* phenotypic class of genes, for example (Zones et al. 2015).

MATLAB code is provided in S.I., which requires as input for each mutant only the vector of Blast/Blosum classes for all SNPs in a mutant, and the likelihood table for Blast/Blosum classes; mapping experiments, consisting of number of recombinant and non-recombinant chromosomes observed relative to a marker, and the physical location of the marker; transcriptional category and the associated likelihood table; and the probability of ‘unsequenceability’ or failure to detect a Ts-lethal lesion by sequencing, which we estimate at ~25% (Tulin and Cross, 2014). Output is the probability of causality for each SNP, as well as probability that the causative mutation escaped detection by sequencing. This code performs well against simulated data: reported probabilities that a SNP is causative are quite accurate, and thresholds can be set with quite high TPR and low FPR (e.g., 95% TPR at 10% FPR; Figure S4).

### Validation of the approach

In Table 1, we report the calculated Bayesian probabilities for causality for candidate SNPs found in genome sequences of mutants in 20 complementation groups. These complementation groups were identified in Tulin and Cross (2014) by single mutant alleles, therefore causative mutation identification was not considered ‘definitive’. Importantly, for this reason none of the mutants or SNPs in Table 2 were used for the training set for Blast/Blosum values or transcriptional pattern.

**Table 2.** Bayesian testing with mutations in 20 genes not in the training set. All 20 *DIV* genes reported in Tulin and Cross (2014) as single alleles not confirmed by reversion testing were analyzed to determine Bayesian probabilities that candidate SNPs in each mutant (full table in S.I.) are causative, based on equations 1a, 2a, 4, and combined tests; triple ‘BLT’ test from equation 5. Tests based on: B: Blast/Blosum values; L: linkage; T: transcriptional pattern; double and triple letters indicate combined tests. In the cases of *div45*, *div46*, *div48*, *div50*, and *div72*, new alleles (‘-2’, ‘-3’) have been isolated in recent mutant screening (Breker, in preparation). Assigned gene model: most likely carrier of causative mutations (annotation information for orthologous *Arabidopsis* gene also provided). Confirmed causality: **YES** indicates isolation of an independent allele (non-complementing, non-recombining) with a lesion in the same gene model. In the case of *div68-1*, we assume the assignment of *DIV68* as a profilin homolog is definitive, even in the absence of a second allele, since *div68-1* mutants have almost no detectable profilin protein by Western blotting (M.Onishi, pers. comm.).

|  |  |  |  |  |  |  |  |  |  |
| --- | --- | --- | --- | --- | --- | --- | --- | --- | --- |
| **div50-2** | 0.90 | 0.38 | 0.84 | 0.90 | 0.98 | 0.84 | **0.98** | **Cre12.g521200/RFC1** | **YES** |
|  | 0.01 | 0.38 | 0.05 | 0.01 | 0.00 | 0.05 | 0.00 |  |  |
| **div51-1** | 0.95 | 0.67 | 0.94 | 0.93 | 0.99 | 0.91 | **0.99** | Cre15.g636300/TFC-B |  |
| **div52-1** | 0.95 | 0.75 | 0.94 | 0.95 | 0.99 | 0.94 | **0.99** | Cre17.g715900/THY2 |  |
| **div53-1** | 0.18 | 0.82 | 0.94 | 0.26 | 0.54 | 0.96 | 0.64 | (Cre17.g744247) |  |
| **div57-1** | 0.58 | 0.90 | 0.94 | 0.98 | 0.99 | 0.98 | **0.96** | Cre06.g278950/Aug4 |  |
| **div60-1** | 0.95 | 0.52 | 0.51 | 0.87 | 0.87 | 0.27 | 0.71 | (Cre01.g039350/P450 reductase) |  |
| **div65-1** | 0.95 | 0.86 | 0.94 | 0.97 | 0.99 | 0.97 | **1.00** | Cre06.g270250/CDC45 |  |
| **div68-1** | 0.82 | 0.26 | 0.76 | 0.85 | 0.97 | 0.78 | **0.98** | Cre10.g427250/Profilin | **YES** |
|  | 0.01 | 0.33 | 0.05 | 0.02 | 0.00 | 0.07 | 0.00 |  |  |
|  | 0.05 | 0.23 | 0.05 | 0.04 | 0.00 | 0.05 | 0.00 |  |  |
| **div70-1** | 0.81 | 0.34 | 0.17 | 0.90 | 0.65 | 0.28 | **0.83** | Cre09.g398650/Cactin |  |
|  | 0.01 | 0.32 | 0.17 | 0.01 | 0.01 | 0.27 | 0.01 |  |  |
|  | 0.06 | 0.22 | 0.17 | 0.04 | 0.05 | 0.18 | 0.04 |  |  |
| **div72-1** | 0.80 | 0.95 | 0.94 | 0.96 | 0.95 | 0.99 | **0.99** | **Cre07.g312350/PolA3,4** | **YES** |
| **div72-2** | 0.95 | 0.75 | 0.94 | 0.95 | 0.99 | 0.94 | **0.99** | **Cre07.g312350/PolA3,4** | **YES** |

Tests in Table 1 used the Blast/Blosum score, linkage, or transcriptional pattern, alone or in all combinations, using the equations above (source data in S.I.). Note that while the initial mutants contained large numbers (tens to hundreds) of coding-sequence-changing mutations, the bulked segregant sequencing strategy (Tulin and Cross 2014), and in some cases additional mapping crosses, whittles these down to a much smaller number, based on the principle that temperature-sensitive segregants WITHOUT some SNP, or temperature-resistant segregants WITH some SNP, eliminate that SNP from consideration for causality (we take this as ‘ground truth’). For each mutant background, all remaining candidate SNPs are tested in the Table. The calculation is carried out by Matlab code with input for each SNP: Blast/Blosum category 1-8, transcriptional category (‘*DIV*-like’ or not), and linkage data (marker location and numbers of recombinant/non-recombinant progeny).

In ~85% of cases, a single SNP is identified as a strongly preferred candidate from each mutant background (calculated probability 0.96 +/- 0.05). In mutants with multiple ‘competing’ SNPs, probability distributions are in general strongly bimodal: one high-probability SNP (usually P>0.9), with other SNPs assigned probabilities of 0.00-0.10. These two clusters most likely represent causative mutations and passengers.

The three criteria (Blast/Blosum, linkage, transcriptional pattern) interact. For the presumed causative mutations (high probability), the single, double and triple tests gave probability estimates of 0.75 +/− 0.24; 0.88 +/− 0.16; 0.96 +/− 0.05 (mean +/− standard deviation); thus combining tests increased the probability estimates. For the presumed passenger mutations, the estimates for single, double and triple tests were 0.15 +/− 0.16; 0.07 +/− 0.13; 0.02 +/− 0.03: combining tests decreased the probability estimates. This is expected if the presumed causative mutations are likely to share all three attributes, while random passenger mutations might fortuitously score high for one but probably not for another one. Thus, the multiple tests interacted to drive divergence in probability estimates between the presumed causative mutations and passengers, resulting in more reliable specific detection. For *gex* mutations, the transcription test is not applicable; nevertheless, the Blast/Blosum and linkage tests interacted in the same way to give high probability identification in most cases.

Critically, new mutant isolation (Breker et al., 2016; Breker et al., in preparation) has resulted in ‘definitive’ determination of 13 causative mutations falling in six genes (*div43*, *div45*, *div46*, *div48*, *div50*, *div72*) (Table 1). This determination is based on the criterion (Tulin and Cross, 2014; see above) that lesions in the same gene model, found in multiple independent isolates in the same complementation group, definitively identify the causative target gene. We also consider the assignment of *DIV68* to the sole profilin homolog in *Chlamydomonas* to be definitive, since the *div68-1* mutation results in almost complete loss of profilin detectable by Western blotting, and also has multiple phenotypes consistent with loss of actin function, as expected for profilin inactivation (M. Onishi, pers. comm.). Thus, 14 mutations, newly determined to be causative, have high calculated Bayesian probabilities (0.93 +/− 0.14).

These results contrast with those for *DIV40. div40-1* contains a single candidate coding-sequence-changing SNP; however, it is a very weak candidate (similar in probability estimates to the collection of known passenger mutations). We now have isolated a second independent allele *div40-2* (defined as allelic since *div40-1* and *div40-2* fail to complement in trans-heterozygous diploids, are both located on the left arm of chromosome 17, and fail to recombine with each other in hundreds of meioses.) However, bulked segregant sequencing of *div40-2* revealed no coding sequence-changing mutation that was uniformly present in Ts− segregants (data not shown), and no mutation at all within hundreds of kilobases of the *div40-1* candidate. Therefore, we suspect that *both* the *div40-1* and *div40-2* causative mutations escaped detection by sequencing, and the single candidate for *div40-1* is indeed a passenger.

This phenomenon, in which multiple independent alleles in a well-defined complementation group fail to share lesions in any one gene model (or have no candidate lesions at all) was observed previously, and was examined with care in the cases of *div14* and *div16* (Tulin and Cross 2014). Four alleles of *div14* were mapped in multiple crosses and by complementation testing to the same position on chromosome 4 (~4 Mb on the physical map), and five alleles of *div16* to chromosome 10 at ~6.5 Mb (data not shown). Bulked segregant sequencing to high coverage (>200X in the cases of one allele each of *div14* and *div16*, and at least 50X coverage of the other seven independent alleles) failed to reveal the causative mutations. We call this phenomenon ‘unsequenceability’, and we estimated previously that ~~25% of temperature-sensitive lethal mutations fall into ‘unsequenceable’ genes (Tulin and Cross 2014). We do not know the explanation for this problem, but it is taken into account in the Bayesian calculation (Appendix). The results on *div40* suggest it may be in this class. This interpretation also supports the value of the Bayesian calculation, since a possible candidate for causation of *div40-1* had a low calculated probability, and was subsequently found to be likely a passenger to an unsequenced true causative mutation.

*div53-1* and *div60-1* mutants are intermediate cases: they contain stronger candidates for causality than *div40-1*, but the candidates are outliers relative to the larger population of presumed causative mutations. The causative mutation in these strains may be atypical with respect to the ‘rules’ followed by the training set. These intermediate cases quantitatively identify mutants for which identification is not strong, and further data are required before relying on the identification for additional experiments.

These results constitute strong validation, since the approach developed with the initial training set of definitive causative mutation generalizes to mutants not in the training set, and in most cases yields a strong preferential identification of a single highly likely causative mutation, even in backgrounds also carrying multiple passenger mutations. The ‘BL’ column shows the power of the Blast/Blosum and linkage tests alone. These tests are sufficient to identify a most-likely candidate in most cases, but it is clear that the orthogonal transcriptional information provides considerable additional resolving power.

## Discussion

### Identification of causative mutations in an unbiased screen

Untargeted mutagenic screens are unbiased, requiring no hypotheses as to which genes might be involved in a given system; this is a clear advantage over targeted approaches such as genome editing. However, to attain reasonable efficiency of isolation of informative mutations, it is necessary to ‘pack’ a large number of mutations into each clone – because most point mutations have little or no phenotypic effect. This means that although high-throughput sequencing can identify (nearly) all mutations in a mutant strain of interest, the problem remains of determining which mutation is causative, among hundreds or thousands of candidates. For any individual mutant, there are methods that will allow absolute certainty as to the causative SNP: high-resolution genetic mapping; sufficient screening to identify multiple independent alleles; isolation of intragenic revertants; rescue by transformation. However, these approaches are impractical for large numbers of mutants.

The Bayesian method converts diverse types of data to the ‘common currency’ of probability. This then allows statement of a quantitative degree of certainty that our identifications are correct. This then provides a rational basis for evaluating the need for further work to confirm the identifications.

### Annotation-independent identification of causative mutations

The problem of identifying causative mutations from a set of candidates is a common one; for example in analyzing cancer genomes, or in population genetics when a QTL is known to be located somewhere in a highly polymorphic haplotype block. In those contexts, it is very difficult to proceed without relying on annotation-based information (e.g., a height QTL with a SNP in a growth-hormone-related gene within the haplotype block). In the experimental context discussed in this paper, genetic methods are sufficiently powerful that annotation-based information can be dispensed with altogether. This is fortunate since first, restriction to annotations largely restricts discovery to things that are already at least partially known; and second, it is not obvious how to assign a quantitative probability value to annotation-based information. The approach above is a uniform Bayesian calculation, which should integrate diverse sorts of information on a quantitatively equal basis.

It is important to note, though, that the power of meiotic mapping to eliminate most SNPs from consideration is essential for success in the present case; the Bayesian discriminators are strong enough to detect one likely positive out of a small number of candidates, but cannot do so from a larger field. This aspect is less applicable in the haplotype block case, and obviously completely unavailable in the cancer genome case. In these situations, additional discriminators are clearly essential.

One class of annotation-based information is required in our approach: parsing of the raw genome sequence into gene models (exons and coding sequence especially). Fortunately, this has been done quite carefully and effectively in the *Chlamydomonas* case (Merchant et al. 2007, Blaby et al., 2014); our detailed examinations of specific issues with the annotation (Cross 2016, Tulin and Cross 2016) have revealed problems with only a small minority of genes.

### Conservation, divergence, gene duplication, and essentiality

A central aspect of this computation is based on the observation that essential genes, identified by ts-lethal single-gene mutations blocking cell proliferation, are much more likely than the *Chlamydomonas* gene set overall to lie in proteins and specific residues conserved in higher plants, and frequently across yeast and animals as well. In contrast, a substantial majority of *Chlamydomonas* genes have either no *Arabidopsis* Blast hit, or only a hit suggestive of a small protein domain. Most of these genes have unknown, possibly algal-specific functions; our results suggest that few of these functions are essential for cell viability, or at the least very seldom are specifically essential for cell cycle progression. Evolution of cell-essential processes is slow.

Isolation of single-gene ts-lethals implies that there is no effective backup in the genome – in particular no gene duplicate retaining substantial functional overlap (since otherwise the lethal phenotype should require at least a double hit). Gene families with presumed orthologous members in *Chlamydomonas* and *Arabidopsis* tend to be single copy in *Chlamydomonas*. In *Arabidopsis*, multiple gene family members are common (Figure 4), a well-known observation substantially due to multiple whole-genome duplications in the higher land plant lineage (Adams and Wendel 2005). It is a commonplace observation in *Arabidopsis* genetics that strong phenotypes frequently require disruption of multiple gene family members. Gene duplication has been proposed to provide genetic ‘robustness’ (Gu et al 2003). In general, the *Chlamydomonas* genome lacks this robustness mechanism: in most cases, mutation of the single *Chlamydomonas* family member has the potential to immediately expose the maximum phenotype (Figure 4). Our results suggest that *Chlamydomonas* has an essential gene set substantially conserved with higher plants, and nearly free of duplicates, supporting its utility as a genetic and cell biological model for the crucially important plant kingdom.

---

**Figure S1.**
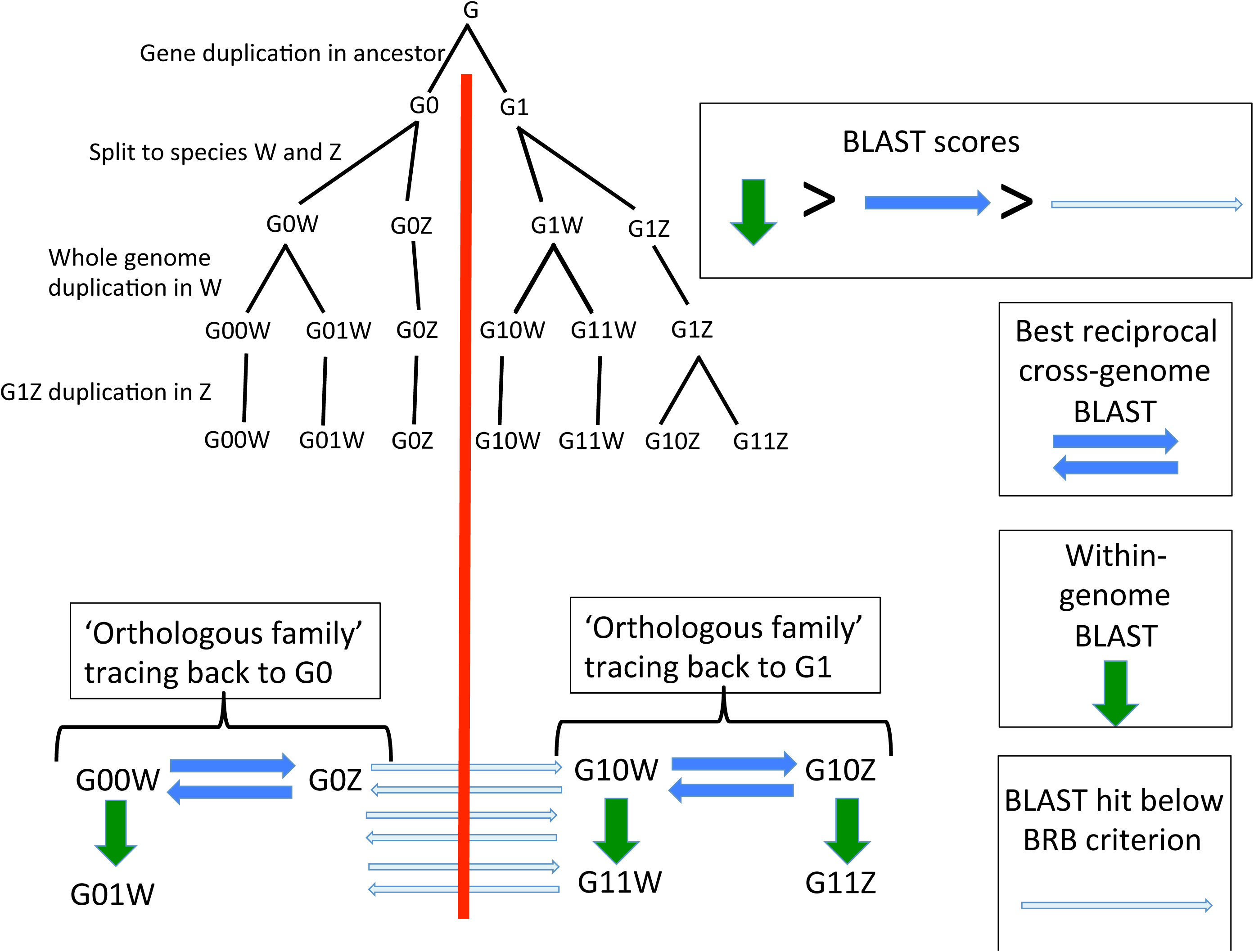
Diagram illustrating how best-reciprocal blast analysis will likely pick out sequences in different genomes originating from a single gene ancestor at the time of the species split.

**Figure S2.**
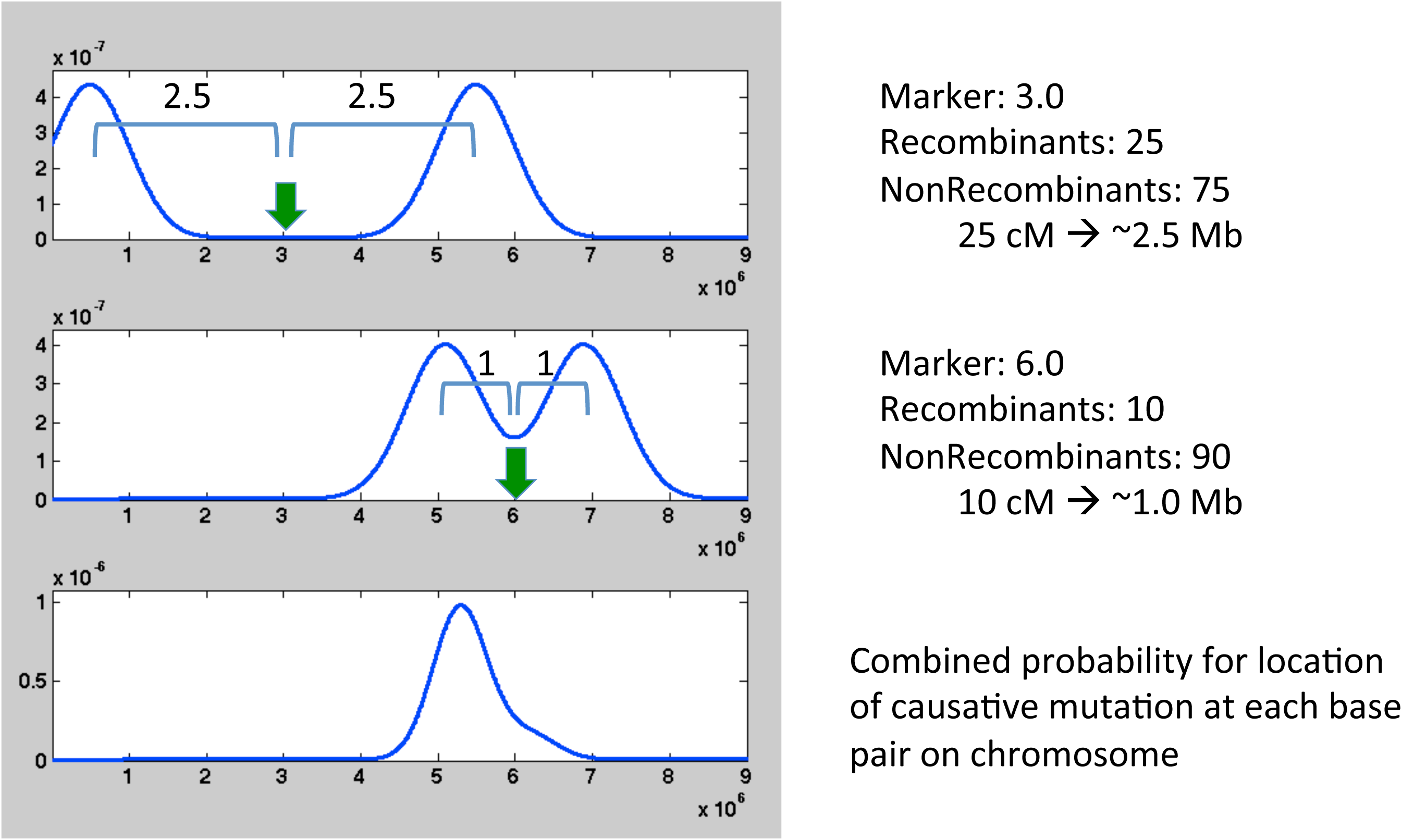
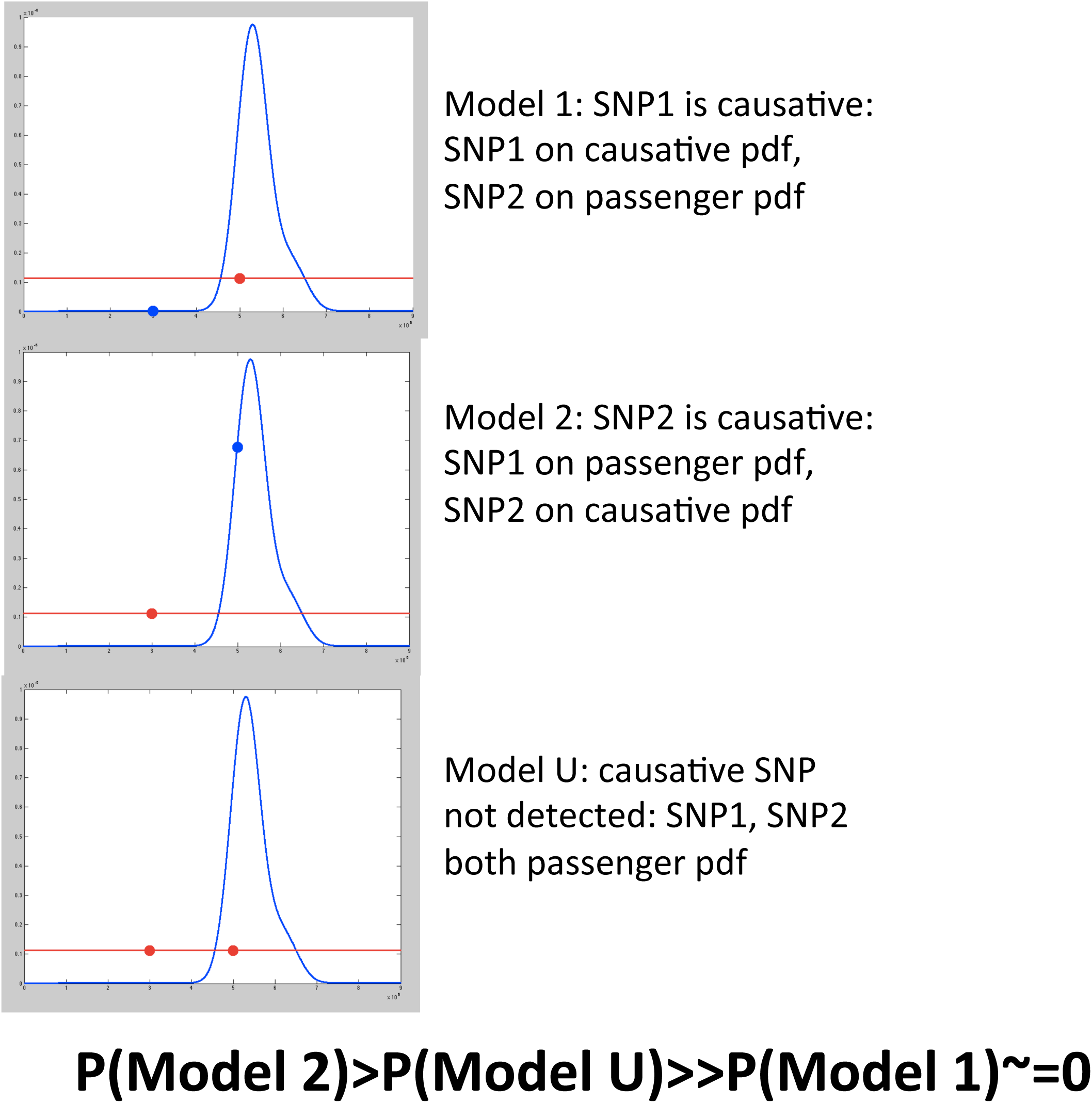
A: Construction of a theoretical probability distribution for location of a causative mutation, based on combining two genetic mapping experiments (top two graphs) in which the ts-lethal mutation was mapped relative to known molecular markers (green arrows). B: if two candidate SNPs are present in this interval, which is known to contain the causative mutation, then the relative probabilities support SNP2 greatly over the alternatives that SNP1 is causative, or that neither SNP is causative.

**Figure S3.**
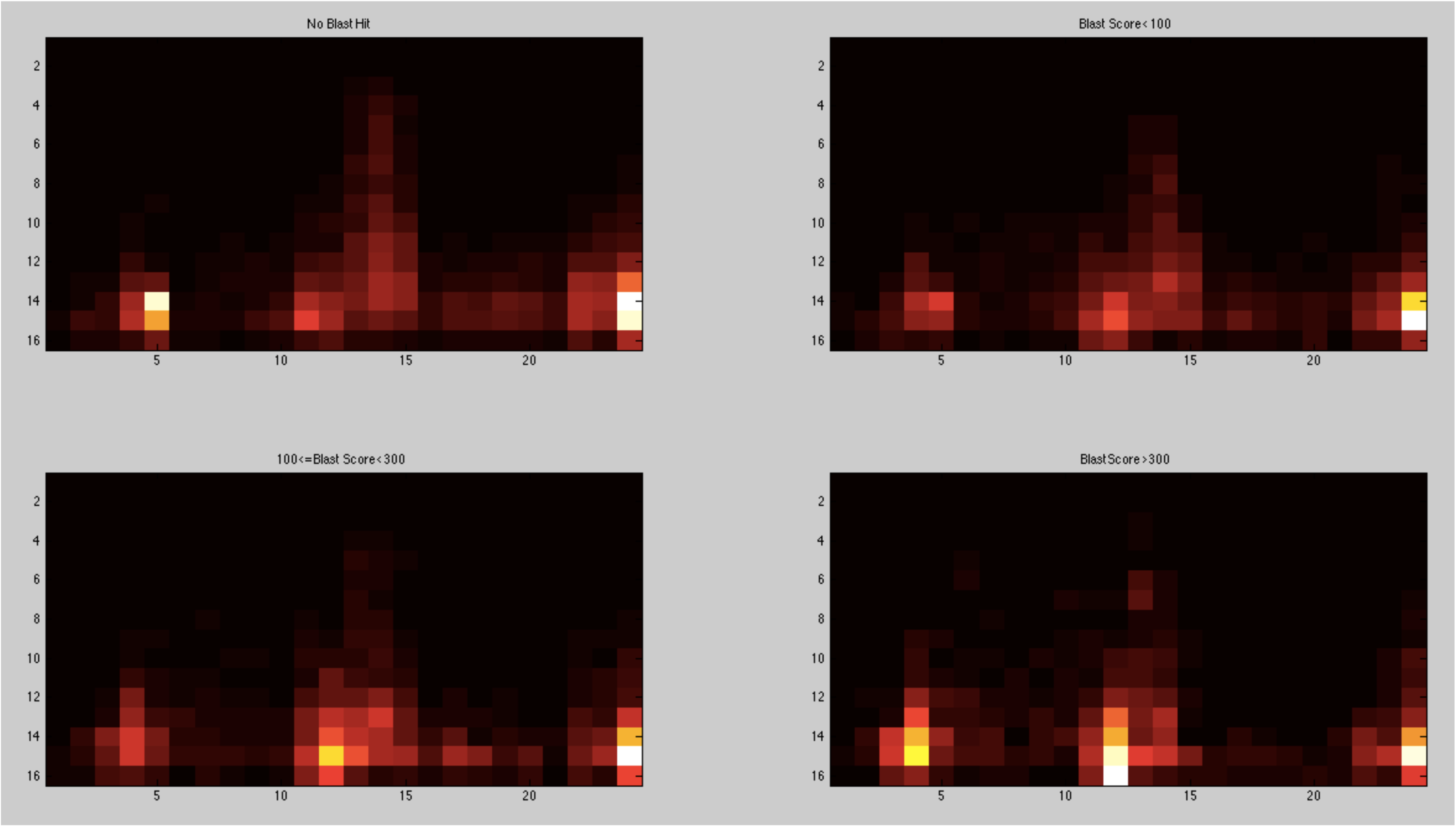
Transcriptional pattern of *Chlamydomonas* genes, segregated by BLAST scores vs. *Arabidopsis*. Overall graphing scheme as in Figure 5, for subsets of genes: Top left: no BLAST hit; 1011/8814 (.11) in DIV-like class; top right: BLAST scores 1-100; 493/3621 (.14); bottom left: BLAST scores 101-300; 545/3564 (0.15); bottom right: BLAST scores >300, 303/1738 (0.17). Overall, transcriptional pattern is largely independent of *Arabidopsis* homology.

**Figure S4.**
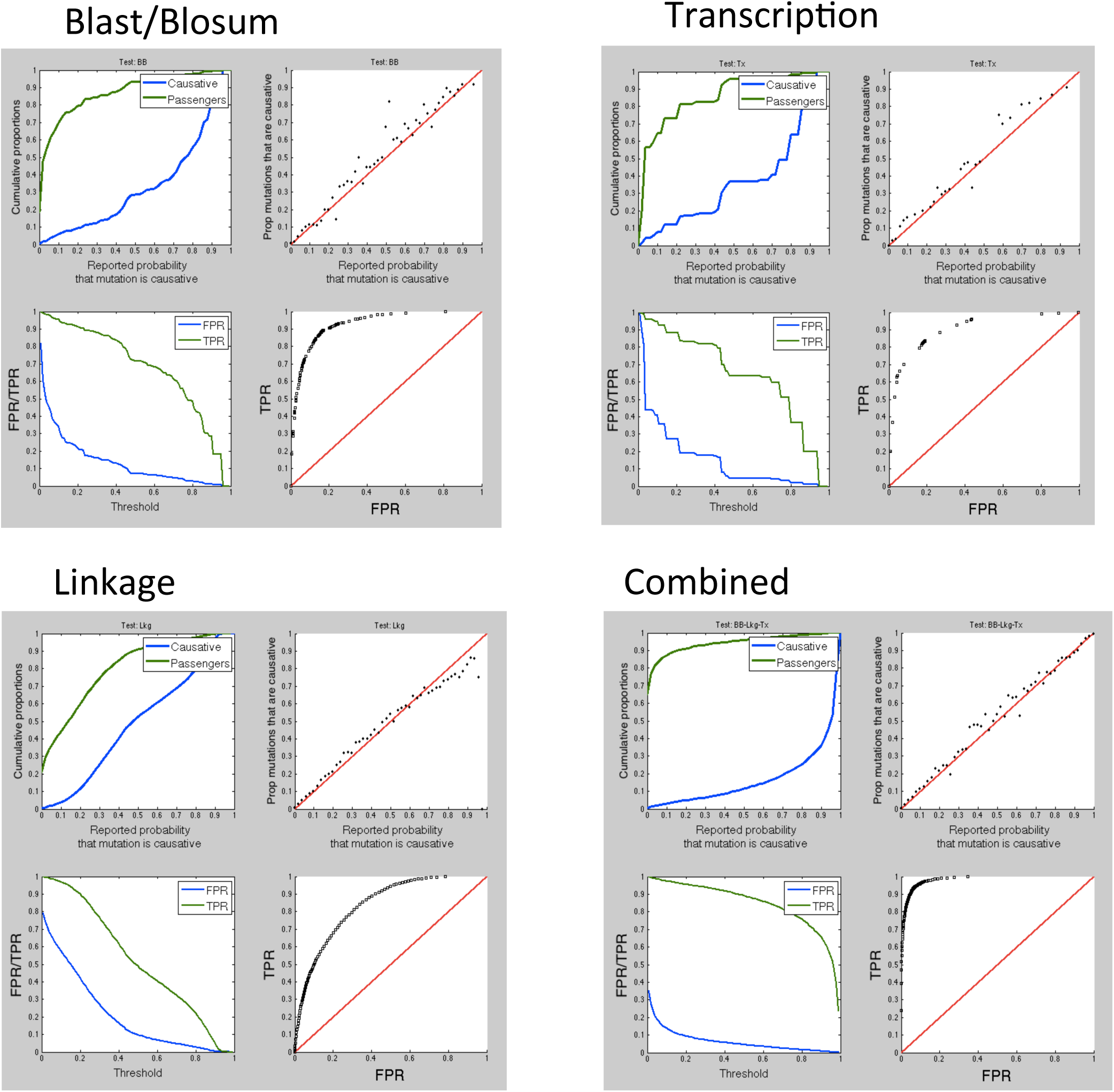
Test of computation with synthetic data. Generated by MATLAB script testMutationDetection.m. 20,000 mutants generated with variable numbers of SNP candidates and a defined position of the causative mutation (which was one of the SNP candidates with probability 0.75; see text). Linkage data was generated explicitly by a simple model assuming strong crossover interference (as observed), and with moderately variable cM/Mb ratios for different mutants. For each SNP the calculated probability that it was causative was recorded, as was the fact of whether it was indeed causative. Distributions of SNPs into Blast/Blosum and Transcription categories followed observed data for passengers and causative mutations. Each set of four graphs: top left: Reported Bayesian probability vs. cumulative proportion of causative (blue) and passenger (green) mutations. Top right: Accuracy: proportion of mutations given some probability of being causative that were in fact causative. The red x=y line is the result for perfect accuracy. Bottom left: false positive rate and true positive rate for different threshold cutoffs. Bottom right: FPR vs. TPR for the data at lower left. Results are shown for the three individual tests (Blast/Blosum, linkage, transcription) and the combined test at lower right.

## Appendix

### detailed development of Bayesian models for Blast/Blosum, linkage, transcription, and combination models

#### Blast/Blosum model

Assume there are N SNPs. Each SNP falls into one of the 8 Blast/Blosum classes. Associated with each class k (k=1:8), there are two likelihoods: Caus(k), the probability that a causative mutation is in class k, and Pass(k), the probability that a passenger mutation is in class k. These values are exactly those plotted in Fig. 2, lower right, derived from the training set.

Then the set of N SNPs is associated with two 1xN vectors C and P:

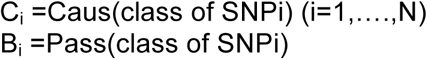

Call U the probability that the causative mutation is *none* of the N candidates (this corresponds to ‘unsequenceability’). (We estimate U at ~25%; Tulin and Cross, 2014).

We assume that either the mutation is unsequenceable (probability U), in which case all of the N SNPs are passengers; or exactly one of the N SNPs is causative; therefore the remaining N-1 are passengers. Call model Mi the model that SNP_i_ is causative. The prior probability of Mi is (1-U)/N, and the probability of MU (unsequenceability) is U. These models are mutually exclusive and exhaustive.

Then the relative probability of S=( S_1_,S_2_…S_n_) (the set of N SNPs) given M_i_ is:

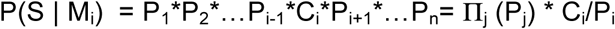

- because in this model, all SNPs are passengers except for SNP_i_. (п_j_ (P_j_) denotes the product of the P_j_’s, j=1 to N)

The relative probability of S given MU (unsequenceable) is:

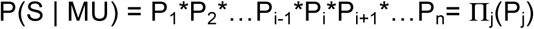

- because in this model, all SNPs are passengers.

These terms are in relative probability units because no probability estimate is provided for having exactly the N SNPs found in S. This probability if expressed would multiply every term and therefore divides out.

Call Q the unsequenceability likelihood ratio U/(1-U).

Bayes’ theorem, given the assumption that exactly one of [M_1_, M_2_,…M_n_, M_U_] must be true, gives:

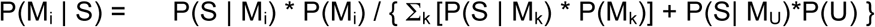

- (where Σ_k_ denotes the sum over k=1 to N)

Substituting:

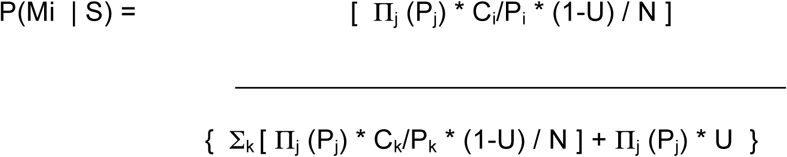

Dividing top and bottom by п_j_ (P_j_), multiplying top and bottom by N/(1-U), and substituting Q for U/(1-U) gives:

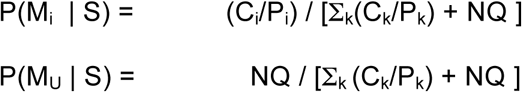

Call B_i_ the likelihood ratio C_i_/P_i_ (the relative prior probability that SNP_i_ is causative relative to the probability that it is a passenger); then

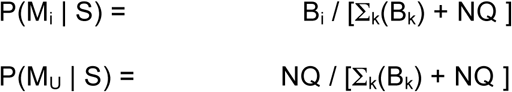

P(M_i_ | S) is the probability that SNP_i_ is causative, given the Blast/Blosum classes of all N candidate SNPs, and the probability U that the causative mutation escaped detection by sequencing.

These probabilities have natural and expected properties. Mutations with high B_i_ (such as severe mutations in conserved residues) have greater probability of being causative (high P(M_i_ | S)). Increasing numbers of candidate mutations (higher N) decreases P(M_i_ | S) – which makes sense since there are many possible candidates to choose from. Mu decreases in probability given the presence of one or more SNPs with high B_i_. This makes sense, since high B_i_ is unlikely; so seeing such a SNP in the collection leads to high likelihood that it is causative. In contrast, if all SNPs have negligible B_i_, the probability of M_U_ approaches 1.

#### Linkage model

In the main text we summarized two methods for meiotic mapping of a Ts-lethal: co-segregation with a SNP marker (in most cases, the SNP suspected of being causative), or segregation compared to a second Ts-lethal with known (or strongly supported) physical map location. These mapping results yield number of recombinant/nonrecombinant chromosomes between a physical marker (the candidate SNP, or the known location of the second Ts-lethal) and the Ts-lethal of interest. The aim is to translate this information into probabilities for physical location of the causative mutation, since then this information can be used as above to discriminate the different models [M_1_, M_2_,…M_n_, M_U_].

To do this, we employed the following quantitative approach. We assume that the physical location of one marker is known (the PCR-detected SNP for the first approach, or the location of the known second mutation in the second approach). We assume an average 10 cM/Mb ratio (Merchant et al 2007, Tulin and Cross 2014), and further assume that the minimum error on the mapping is 5 cM~=0.5 Mb, based on the largest coldspot we have detected, as well as apparent mapping errors over a large number of such experiments (Tulin and Cross 2014). In almost all experiments, sufficient meioses are tested that this uncertainty (rather than sampling error) is the main source of error.

We then suppose that the probability density for mutant location is approximately normal, with mean at the exact estimated location. The standard deviation is estimated by finding 95% confidence limits for the true recombination rate, given the observed numbers of recombinants and nonrecombinants. For a normal distribution, these 95% confidence limits will be separated by ~4 standard deviations. The theoretical standard deviation is then set as the minimum of this distance/4, and 0.5 Mb (to account for uncertainty about the uniformity of the 10 cM/Mb genomic average; see above). Mapping is bidirectional (e.g. 10 cM from a marker could be 10 cM to the left or the right), so we construct both normals, sum them and renormalize to make the area 1. In cases where the distribution is terminated (the end of the chromosome, or a the end of a region of uniformity in bulked segregant sequencing analysis), we truncate the distribution and renormalize. This also means that probability of causality on unlinked chromosomes is set to zero. Note that presence of the causative mutation on this limited region is set as ‘ground truth’ in this approach. This is reasonable, since linkage to a chromosome is generally established already to extremely high probability.

If there are multiple such mapping experiments, the probability densities are multiplied at each point and renormalized to make a single model. Thus, for example, a bimodal distribution of likely locations is converted by this multiplication to an essentially unimodal one, by finding cosegregation of Ts-lethality with a SNP near the center of one of the bimodal peaks.

An illustration of generation of such a probability density function is shown in Fig. S2A, assuming one experiment showing 25/100 recombinants of a marker at 3.0 Mb with Ts-, and another showing 10/100 recombinants of a marker at 5.0 Mb with Ts-. The two mapping experiments (top and middle) are bimodal because distance is approximately known (using 10 cM/Mb conversion) but direction is not. The two combined (bottom) strongly favor a location of the Ts- lesion at ~5.5 to be consistent with both mapping experiments. All curves have total area 1. Call this function f, where f(location)=probability of the causative mutation being at the location.

f is in units of (probability/basepair), and is nonzero only over the interval known (as ground-truth) to contain the Ts-lethal mutation.

Thus, the probability density for location of the causative SNP should be just f.

In contrast, the probability density for a passenger SNP should be uniform over the ‘ground-truth’ interval, zero elsewhere, because presence somewhere in this interval is required for it to be in consideration. Thus this probability is identically 1/(length of ground-truth interval) for all candidates (all positions equally likely). Call the length of this interval G; so probability for passengers at each basepair in the interval is 1/G (units of probability/basepair, the same units as for f).

Call L_i_ the location of SNPi (L is the vector over the N SNPs). Then:

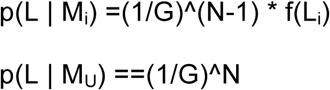

Rescale probability density for passenger and causative SNPs by multiplying by G; then the relative pdf of each passenger is 1, and that of the causative SNP is G*f. Call G*f function g.

Then:

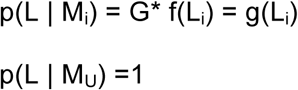

- where p is proportional to probability density.

Prior probabilities of M_i_’s and M_U_ are as before.

Then Bayes’ theorem yields:

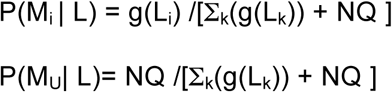

(Scaled probability density above is converted to probability here, because all terms are in the same units of relative probability density).

Figure S2B shows placement of two SNPs on the probability curves according to models [M_1_, M_2_, M_u_] (only two candidate SNPs in this example). In this case M2 is most likely (product of probabilities for the two SNPs is highest for this model).

#### Combined model

Note that clearly, location of a SNP on the chromosome is independent of the Blast/Blosum characteristics of the SNP. This means that probabilities multiply. So:

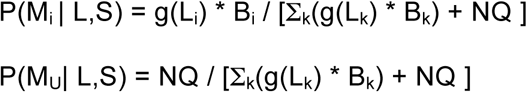

#### Transcription

For the *DIV* subclass, we can also integrate the probability that SNP_i_ is causative, based on whether its containing gene follows the transcriptional pattern of most of these genes (since transcriptional pattern is essentially independent of the other classifications). Using the set of strongly identified *DIVs*, we have a simple 2X2 classification table: *DIV* gene vs. all genes, S/M transcription pattern vs. different pattern (Figure 5). If Ti is the likelihood ratio for the gene model containing SNP_i_ relative to transcription pattern, based on the training set of definitive *DIVs* vs. all genes (Figure 5), we can integrate this information into the calculation:

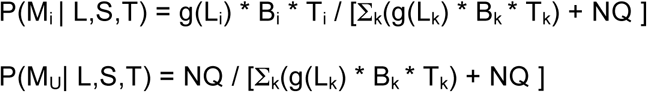

To confirm that the model is correctly generated and the code calculating probabilities accurately, we generated 20,000 ‘mutants’ in silico (Figure S4), aiming for a reasonable simulation of the observed distributions, and number and scale of typical linkage experiments. High accuracy, sensitivity and selectivity were observed, verifying the correctness of the calculations.

## REFERENCES

Adams KL, Wendel JF. 2005 Polyploidy and genome evolution in plants. Curr Opin Plant Biol. 2005 Apr;8(2):135–41.

Blaby IK, Blaby-Haas CE, Tourasse N, Hom EF, Lopez D, Aksoy M, Grossman A, Umen J, Dutcher S, Porter M, King S, Witman GB, Stanke M, Harris EH, Goodstein D, Grimwood J, Schmutz J, Vallon O, Merchant SS, Prochnik S. The Chlamydomonas genome project: a decade on. Trends Plant Sci. 2014 Oct;19(10):672–80.

Botstein D, Fink GR. 2011Yeast: an experimental organism for 21st Century biology. Genetics. 2011 Nov;189(3):695–704.

Breker, M, Lieberman, K, Tulin, F, Cross, FR. High-throughput robotically assisted isolation of temperature-sensitive lethal mutants in Chlamydomonas reinhardtii. Journal of Visualized Experiments, in press.

Cross FR. Tying Down Loose Ends in the Chlamydomonas Genome: Functional Significance of Abundant Upstream Open Reading Frames. G3 (Bethesda). 2015 Dec 23;6(2):435–46.

Cross FR, Buchler NE, Skotheim JM. 2011. Evolution of networks and sequences in eukaryotic cell cycle control. Philos Trans R Soc Lond B Biol Sci. 2011 Dec 27;366(1584):3532–44.

Gu ZL, Steinmetz LM, Gu X, Scharfe C, Davis RW, Li WH. Role of duplicate genes in genetic robustness against null mutations. Nature. 2003;421: 63–66.

Henikoff S, Henikoff JG. Performance evaluation of amino acid substitution matrices. Proteins. 1993 Sep;17(1):49–61.

Medina EM, Turner JJ, Gordân R, Skotheim JM, Buchler NE. Punctuated evolution and transitional hybrid network in an ancestral cell cycle of fungi. Elife. 2016 May 10;5.

Merchant SS, Prochnik SE, Vallon O, Harris EH, Karpowicz SJ, Witman GB, Terry A, Salamov A, Fritz-Laylin LK, Maréchal-Drouard L, Marshall WF, Qu LH, Nelson DR, Sanderfoot AA, Spalding MH, Kapitonov VV, Ren Q, Ferris P, Lindquist E, Shapiro H, Lucas SM, Grimwood J, Schmutz J, Cardol P, Cerutti H, Chanfreau G, Chen CL, Cognat V, Croft MT, Dent R, Dutcher S, Fernández E, Fukuzawa H, González-Ballester D, González-Halphen D, Hallmann A, Hanikenne M, Hippler M, Inwood W, Jabbari K, Kalanon M, Kuras R, Lefebvre PA, Lemaire SD, Lobanov AV, Lohr M, Manuell A, Meier I, Mets L, Mittag M, Mittelmeier T, Moroney JV, Moseley J, Napoli C, Nedelcu AM, Niyogi K, Novoselov SV, Paulsen IT, Pazour G, Purton S, Ral JP, Riaño-Pachón DM, Riekhof W, Rymarquis L, Schroda M, Stern D, Umen J, Willows R, Wilson N, Zimmer SL, Allmer J, Balk J, Bisova K, Chen CJ, Elias M, Gendler K, Hauser C, Lamb MR, Ledford H, Long JC, Minagawa J, Page MD, Pan J, Pootakham W, Roje S, Rose A, Stahlberg E, Terauchi AM, Yang P, Ball S, Bowler C, Dieckmann CL, Gladyshev VN, Green P, Jorgensen R, Mayfield S, Mueller-Roeber B, Rajamani S, Sayre RT, Brokstein P, Dubchak I, Goodstein D, Hornick L, Huang YW, Jhaveri J, Luo Y, Martínez D, Ngau WC, Otillar B, Poliakov A, Porter A, Szajkowski L, Werner G, Zhou K, Grigoriev IV, Rokhsar DS, Grossman AR. et al. 2007 The Chlamydomonas genome reveals the evolution of key animal and plant functions. Science. 2007 Oct 12;318(5848):245–50.

Onishi M, Pringle JR, Cross FR. Evidence That an Unconventional Actin Can Provide Essential F-Actin Function and That a Surveillance System Monitors F-Actin Integrity in Chlamydomonas. Genetics. 2016 Mar;202(3):977–96.

Remm M, Storm CE, Sonnhammer EL. Automatic clustering of orthologs and in-paralogs from pairwise species comparisons. J Mol Biol. 2001 Dec 14;314(5): 1041–52.

Rogozin IB, Basu MK, Csürös M, Koonin EV. 2009 Analysis of rare genomic changes does not support the unikont-bikont phylogeny and suggests cyanobacterial symbiosis as the point of primary radiation of eukaryotes. Genome Biol Evol. 2009 May 25;1:99–113.

Tulin F, Cross FR 2014. A microbial avenue to cell cycle control in the plant superkingdom. Plant Cell. 2014 Oct;26(10):4019–38.

Tulin F, Cross FR. Cyclin-Dependent Kinase Regulation of Diurnal Transcription in Chlamydomonas. Plant Cell. 2015 Oct;27(10):2727–42.

Tulin F, Cross FR. Patching Holes in the Chlamydomonas Genome. G3 (Bethesda) 2016 Jul; 6(7): 1899–1910.

Yampolsky LY, Stoltzfus A. The exchangeability of amino acids in proteins. Genetics. 2005 Aug;170(4):1459–72.

Zones JM, Blaby IK, Merchant SS, Umen JG. High-Resolution Profiling of a Synchronized Diurnal Transcriptome from Chlamydomonas reinhardtii Reveals Continuous Cell and Metabolic Differentiation. Plant Cell. 2015 Oct;27(10):2743–69.

